# Contextual Prediction Errors Reorganize Episodic Memories in Time

**DOI:** 10.1101/2020.08.24.265132

**Authors:** Fahd Yazin, Moumita Das, Arpan Banerjee, Dipanjan Roy

## Abstract

Episodic memories are contextual experiences ordered in time. This is underpinned by associative binding between events within the same contexts. The role of prediction errors in strengthening declarative memory is well established but has not been investigated in the time dimension of complex episodic memories. Here we used 3-day movie viewing paradigm to test the hypothesis that contextual prediction errors leads to temporal organization of sequential memory processing. Our main findings uncover that prediction errors lead to changes in temporal organization of events, secondly, new unexpected sequences show as high accuracy as control sequences viewed repeatedly, and these effects are specifically due to prediction errors, and not novel associations. A drift-diffusion modelling further revealed a lower decision threshold for the newer, unexpected sequences compared to older sequences reflected by their faster recall leads to reorganization of episodes in time. Moreover, we found individual decision threshold could significantly predict their relative speed of sequence memory recall. Taking together our results suggest a temporally distinct role for prediction errors in ordering sequences of events in episodic memory.

## Introduction

Imagine while entering the office, you see your favorite actor sitting in your chair, much to your surprise. The memory of such an event would be harder to forget than others. The substantial memory consolidation of this example event, due to the low expectation of such events occurring within the given context displays the dependency of our day-to-day memories on the underlying contextual^1, 2, 3, 4, 5^ and predictive processes^6, 7, 8, 9, 10^. Episodes or events being the canonical components of episodic memory^11^, are not only marked by a clear beginning and an end, but also relatable to each other temporally^11^. As such, our memories are organized sequentially in contexts that evolves in time. But whether and how unpredicted events can affect this temporal code of our experienced memories is something that surprisingly remains largely unexplored.

Predictions are a hallmark of episodic memory recall^6, 7,10^. Therefore, whenever a context is re-experienced, the sequence of episodes is automatically predicted^6, 9, 10, 12^. Prediction errors resulting from the violation of these predictions have been shown to influence declarative memory by strengthening incidental encoding^13^, semantic memory acquisition^14^, paired association learning^15, 16^ and by playing a role in reconsolidation^12^. Despite its wide effects on declarative memory, its role in episodic memories are only uncovering now.

A core property of episodic memory is the sequential arrangement of the events as and how they occurred in their respective contexts. A context in its simplest form is described as any aspect of the episode that binds its constituent elements together, be it spatial, temporal or conceptual^2, 3, 4, 5, 11^. Daily life involves numerous instances where multiple different event instances share the same context. The Temporal Context Model (TCM)^1, 17^ of memory posits that such memories sharing the same temporal context are encoded separately, creating source confusion during memory recall. Memories of items shared in same context have been observed to be weakened^18^. It can also be explained by an important line of work termed as reconsolidation^4, 19, 20, 21, 22^, which posits the older memories would get updated which is thought to be mediated by prediction error^23, 24, 12^ and is demonstrated by an asymmetric intrusion of memories in the same context during recall. However, a recent theory^2^ which builds on and unites a lot of theoretical frameworks of memory, proposed contextual binding as a unified mechanism with the hippocampus playing a central role in item and context binding. In addition to hippocampal associative learning mediating context representation, this theory also posits that forgetting occurs mainly due to contextual interference from shared memories. The hippocampus, interestingly is also predictive in nature^10, 25^ and is sensitive to prediction mismatches^26^. This sets up an intriguing questions on how interactions between these two properties affect episodic memories.

In the present study, we test the hypothesis that the contextual prediction errors would fundamentally alter the memorized sequence of events. Specifically, sequences that are predicted but not experienced in a context would be weakened, and the new sequence that was seen instead would be strengthened, as a whole. From the perspective of predictive coding^27, 28^ new surprising information drives associative learning and the ensuing sequential order would be strengthened over older encoded sequential information, so as to minimize the errors in the future. Our key finding is that contextual prediction errors strengthens the newer memory sequences in time while weakening the order of previously encoded sequences, thereby reorganizing encoded temporal memories. This enhanced performance, reflected by faster reaction times, on subsequent modelling showed that it results from lower decision threshold while remembering, signifying a more automatic response for the newer sequences. Critically, even the reexposure of mispredicted segments in an event later on did not exempt it from getting weakened while recalling. Collectively our findings reveal how prediction errors play a key role in determining how episodic memories are organized in time.

## Results

### Formation of contextual priors and subsequent prediction errors

To systematically validate our hypothesis, we employed a 3-day paradigm with naturalistic movies strategically edited into contextually different events containing multiple segments (Fig. 1a). Since one of our main goals was to understand the crucial relationship between prediction error and the evidence accumulation process for the subsequent memories during sequence recall, we devised a way to measure the strength of individual temporal memories for the movie events. After watching the movies on *Day1*, participants saw an incongruent version of the movies with different but contextually fitting segments added or replaced onto the original event during *Day2* with a sequence memory test conducted on *Day3*. The test required participants to choose the correct temporal order of adjacent segments within the same event (Fig. 1c). Moreover, we had two conditions in which participants watched the movies – *Substitution* condition (Fig. 1b, top), where a segment is replaced by another segment having different content (on *Day2*), but fitting with the context, and *Addition* condition (Fig. 1b, bottom), where after viewing the *New* segment the *Old* segment was viewed again. This second condition allowed us to understand the relationship between prediction error and reactivation, a property of memory which strengthens them.

**Fig. 1.**
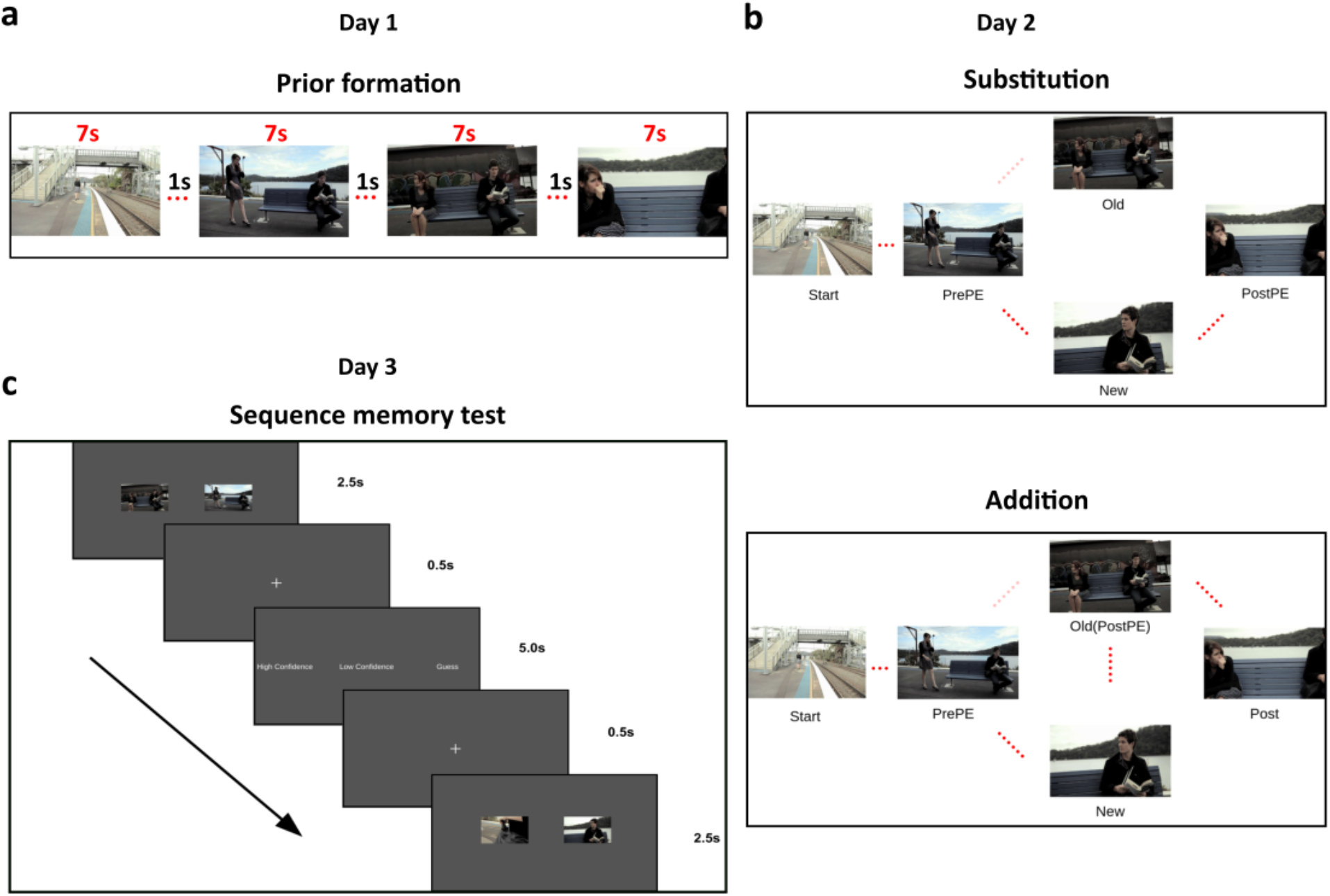
Experiment paradigm. Participants watched two movies (divided into several different events having multiple segments) on *Day1*. The following day (*Day2*), they saw the same movies in either of two conditions - *Substitution* and *Addition*. *Substitution* had another contextually fitting segment substituting a prior encoded segment, while in *Addition* the omitted segment is viewed again (after the prediction error). A sequence memory task (2-AFC) of adjacent segments tested participants’ temporal order memory for each event on *Day3*. **a**, Example of an event seen on *Day1*. Each segment is 7s with a 1s blank screen in between (not shown). **b**, *Day2* conditions. *Substitution* (top) where participants were predicting a segment (*Old*) seen the previous day (faded red dots) while actual segment (*New*) is a different one which fits with the context. *Addition* (bottom) which has the *Old* (predicted) segment re-experienced (hence reactivated) after the prediction error is induced by the *New* segment. *PrePE-Old* temporal sequence memory is taken as Old sequence and *PrePE-New* temporal sequence is taken as New sequence. **c**, Schematic of *Day3* Sequence memory test block. Each sequence was shown by displaying representative screenshots of those segments involved (in both normal and reverse order). Participants had to choose the correct order of the segments in the order they saw in the movie.

### Prediction errors reorganize temporal episodic memories

We hypothesized that the association strength of the inaccurately predicted Old sequences would be weaker compared to the New sequences because of the prediction error. Subsequently, we compared the sequential order memory between *PrePE* segment and *Old* segment with the *PrePE* segment and the *New* segment. Indeed, there was a significant difference in percentage accuracy between recalling the Old (Mean = 47.69, SE = 3.04) and New sequence (Mean = 58.76, SE = 3.05) in *Substitution* condition (*t_(23)_*= 3.416, *p* = .002, 95% CI [4.36, 17.77], BF = 16.60, *d* = .74) (Fig. 2a). There was a significant decrease in reaction time for the New sequence (Mean RT = 1517ms, SE = 34ms) compared to Old sequence (Mean RT = 1600ms, SE = 44ms) *t_(23)_ =* −2.42, p = .02, 95% CI [−151.8, −12.13], BF = 2.39, *d* = .42 (Fig. 2a).

**Fig. 2.**
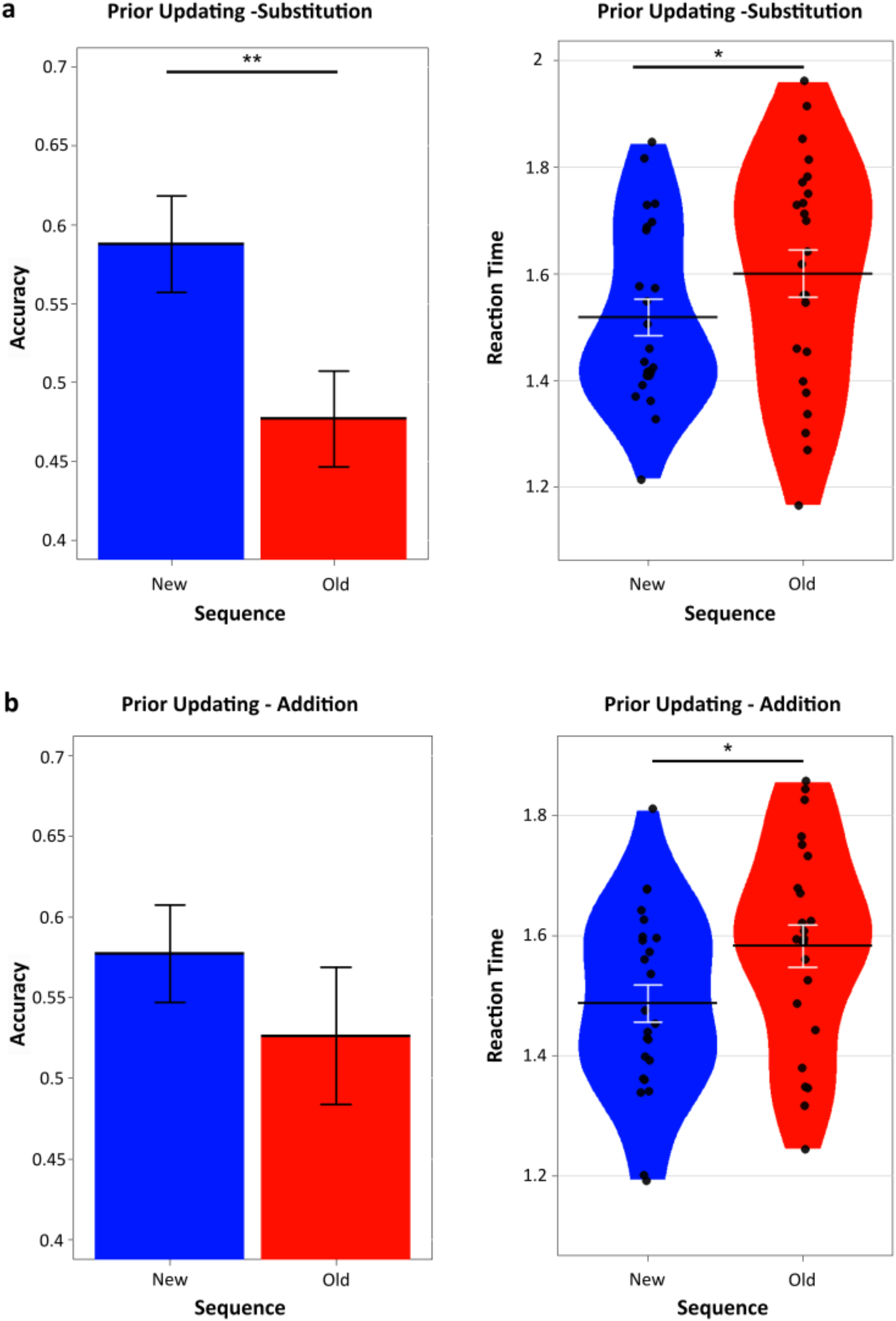
Prediction errors affect temporal order of the memory by updating prior encoded sequences. **a**, Accuracy and reaction times for *Substitution* condition showing significantly greater memory performance for the New sequences over Old sequences. **b**, In *Addition* condition, reactivation of the initially omitted memories leading to re-experiencing the segments again shows no significance in memory accuracy between the sequences but Old sequences show significantly slower RTs. Dots represent participants’ individual performance (*n* = 24). Error bars denote s.e.m.

Mental representation of event sequences helps predict temporal causality of the segments. A PrePE segment would be predictive of the upcoming Old segments after initial viewing. After a PE occurred however, the same PrePE segment was more predictive of the New segment. The effects of prediction errors on the accessibility of the memory sequences is reflected by this result. In other words, the New sequences are always recalled faster compared to the Old ones.

### Slower recall of reactivated memories with prediction errors

Next, we hypothesized that the segments omitted in *Substitution* if re-experienced again, would result in stronger memory activation of those segments. That is, if the mispredicted segment is re-experienced, its subsequent reactivation would strengthen its sequential memory. The *Addition* condition was used to test out this specific hypothesis. This allowed us to tease out the specific effect of PE by comparing New and Old Sequences by its interaction with memory reactivation. Furthermore, this also allowed us to control for a potential temporal recency confound in *Substitution* condition. In other words, whether the results observed in Substitution is due to the fact that New sequences had segments seen on Day2 compared to Old sequences having segments from Day1, potentially giving a recency advantage. We found that the group differences in mean accuracy of New (Mean = 57.71, SE = 3.02) and Old (Mean = 52.62, SE = 4.25) sequences did not differ significantly *t*_*(23)*_ = 0.906, *p* = .37, 95% CI [−6.5, 16.69], BF = 0.31, *d = .28* (Fig. 2b). This is not surprising given that in *Addition* condition re-encountering of the *Old* segment after the prediction error occurs, in contrast to *Substitution* condition where the segment was omitted. Reaction times however, showed a significant difference (New Sequence Mean RT = 1486ms, SE = 31ms, Old Sequence Mean RT = 1582ms, SE = 35ms), similar to the *Substitution* condition *t*_*(23)*_ = −2.43, *p* = .02, 95% CI [−177, −14.32], BF = 2.41, *d = .58)* (Fig. 2b). These results suggest a specific effect of PEs on the New memory sequences reflected by their faster RTs with concomitant slowing of Old memory sequences, even if reactivated.

### A one-shot learning observed in the New sequences

Recent studies have put forth a one-shot encoding property of PEs^16^. We wondered whether this holds true in temporally extended naturalistic memories as well. We compared the Control sequences, (*Start*-*PrePE*) so named since they are seen repeatedly on *Day1* and *Day2* without any changes or violations with the New (PE) sequences. This enabled us to contrast how much of learning did PEs enhance in the memory accuracy, compared to a sequence experienced twice.

Remarkably, in *Substitution* there was no significant difference (*t*_*(23)*_ = 0.83, *p* = .41, 95% CI [−0.054, 0.126], *d* = 0.23, BF = 0.29) between New (Mean = 58.76, SE = .03) and Control sequences (Mean = 62.38, SE = .03). (Fig 3a). This indicates that learning of event sequences occurring through repetitive encoding and one-shot encoding can have comparable accuracies if a prediction error was produced in learning of the latter.

**Fig. 3.**
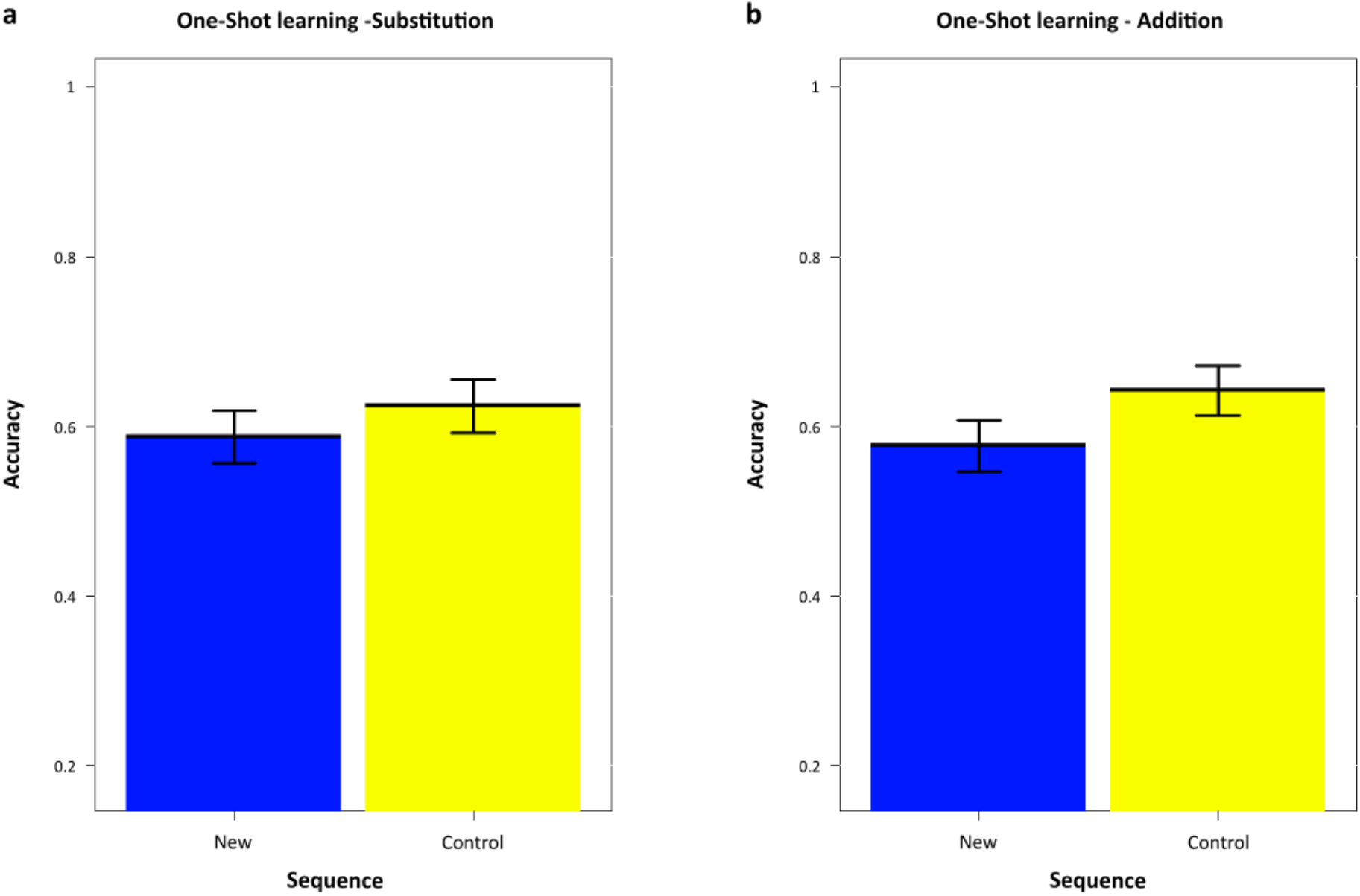
One-shot learning exhibited by New sequences. Similar accuracy performance in sequence memory recall between New (blue) and Control (yellow) sequences in **a**, Substitution and **b**, Addition suggesting a one-shot style learning. Control sequence is the sequence of *Start-PrePE,* which was unchanged for both conditions and days. Dots represent participants’ individual performance (*n* = 24). Error bars denote s.e.m.

This effect of PEs on the memories was noted in *Addition* condition as well with similar memory accuracies (*t*_*(23)*_ = 1.54, *p* = .13, 95% CI [−0.022, 0.152], *d* = 0.44, BF = 0.60) between New (Mean = 57.71, SE = .03) and Control sequences (Mean = 64.22, SE = .029) (Fig. 3b). Firstly, this shows the effect is present independent of conditions and secondly it solidifies the importance of prediction errors in driving one-shot episodic sequence learning.

### The effects on temporal order memory are specifically because of PEs and not due to novel associations

In order to truly verify that the memory effects are due to PEs but no other factors like novel associations, we compared the New sequence (*PrePE*-*New*) with the subsequent sequence, *PostPE*-*New*, which was termed as Novel sequence. The reasoning being, since both these memory associations are encoded on *Day2*, unless there was a specific effect of PEs, both would justifiably have similar memory encoding. Thus, we hypothesized since the memory strengthening can only be due to PEs, the New sequences would have significantly better memories over novel associations.

In *Substitution*, there was a significant difference (*t*_*(23)*_ = 2.12, *p* < .05, 95% CI [0.002, 0.214], BF = 1.41, *d* = .67) in accuracy between New sequence, (Mean = 58.76, SE = .03) compared to the Novel sequence (Mean = 47.88, SE = .03) (Fig. 4a). This difference was significant in response times as well (*t*_*(23)*_ = −3.07, *p* = 0.0054, 95% CI [−0.136, −0.026], BF = 8.18, *d* = .44) between *New* Sequence (Mean RT = 1517ms, SE = 34ms) and the Novel Sequence (Mean RT = 1600ms, SE = 41ms) (Fig. 4a). This shows that in *Substitution*, even though both the sequences were seen on the same day, there is a stark difference in memory performance specifically due to PEs and not seen in novel sequences. Next, we sought to replicate this in the *Addition* condition.

**Fig. 4.**
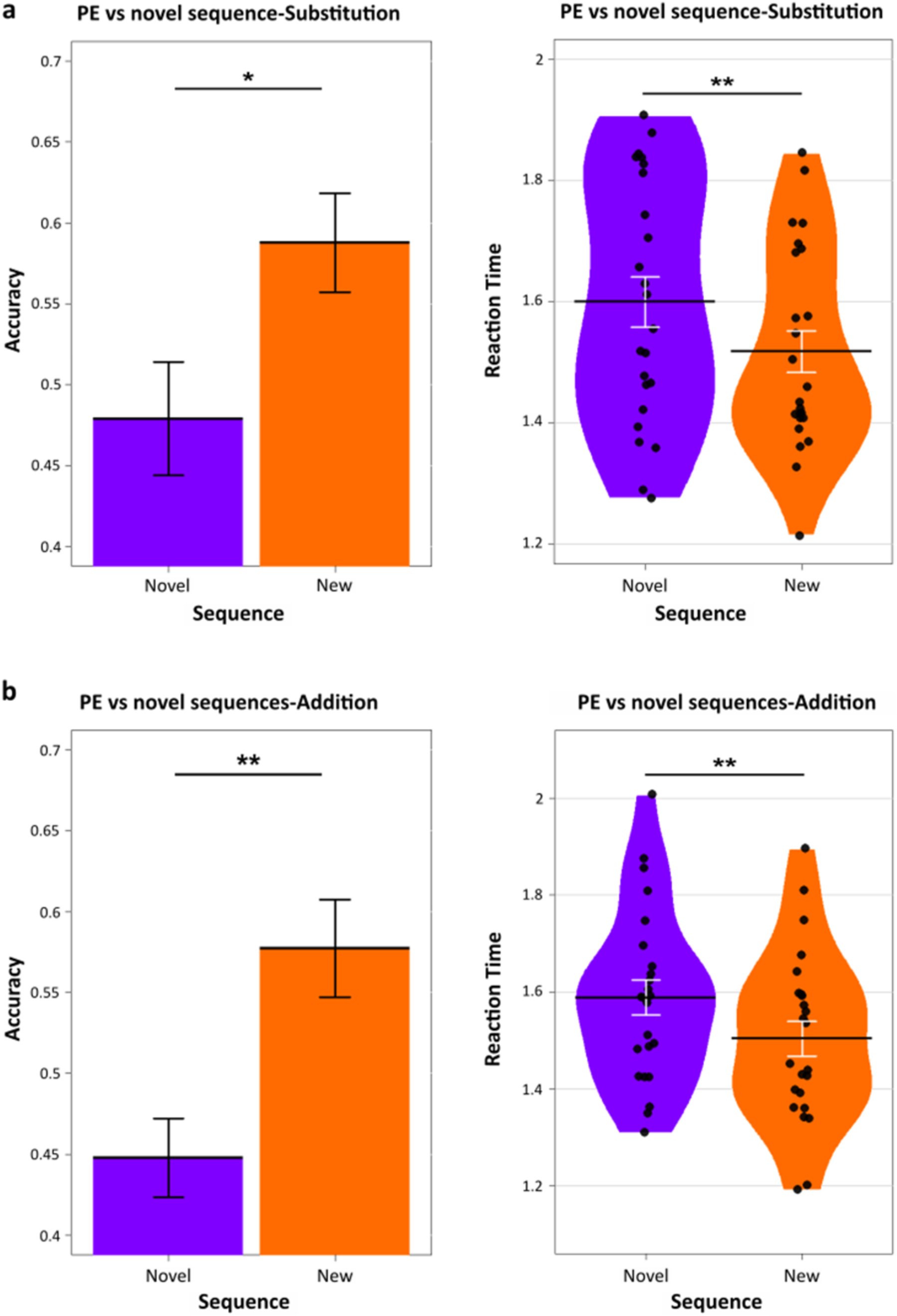
Effects are due to Prediction Errors and not novel associations. New sequences were compared to novel sequences, which was the *New-PostPE* sequence **a**, Accuracy and Reaction times in sequence memory of New (PE) sequence (Orange) and novel sequence (Purple) in *Substitution* condition denoting better performance in New sequences but not in novel sequences. **b**, Accuracy and Reaction times in sequence memory of New (PE) sequence (Orange) and Novel sequence (Purple) in *Addition* condition. Note that the novel sequence here is *New-PostPE* sequence of which the *PostPE* is the segment that was originally omitted but seen again. Dots represent individual participant performance. Error bars represent s.e.m.

We hypothesized that similar patterns would be present in *Addition* as well. A significant difference (*t*_*(23)*_ = 3.29, *p* = 0.0031, 95% CI [0.048, 0.210], BF = 12.8, *d* = .96) was observed in accuracy between New sequence (Mean = 57.71, SE = .03) compared to the Novel sequence here as well (Mean = 44.77, SE = .02) (Fig. 4b). Response times demonstrated significant differences as well (*t_(23)_* = −2.82, *p* = 0.0096, 95% CI [−0.175, −0.027], BF = 5.00, *d* = .61) (*New* Sequence Mean RT = 1486ms, SE = 31ms, Novel Sequence Mean RT= 1588ms, SE = 36ms) (Fig. 4b). This conclusively shows the memory strengthening effects are exclusive when strong prior expectations are violated, but not when mere associations (without any explicit prediction errors) are formed.

### Prediction errors reduce decision threshold during sequence recall of newer memories

The main model allowed both drift rate and boundary parameters to vary with Stimulus (*Old* and *New* memory sequences). This allowed us to compare which of the two parameters demonstrated the empirical effect of PEs, which is the reaction time of the participant in choosing the temporal order between two segments. Drift rate also varied across Confidence (High Confidence and Low Confidence) (Supplementary Table 1).

Thus our key question from a modelling perspective was to test whether faster memory retrieval performance is dependent on lower decision threshold or faster drift rates. The best fitting model (Supplementary Fig. 4) had both drift rates and boundary parameter allowed to vary with the Stimulus, that is Old and New sequences. Drift rate was also allowed to vary with Confidence measures for each trial response. The HDDM model reliably accounted for reaction times. Moreover, a posterior predictive check was conducted to assess model performance (by generating 100 datasets from the model) which could predict the observed RTs (Fig. 6a, Fig. 6c).

Significant differences between the Boundary parameter *a* of Old and New sequences explained the empirical data (Boundary_*Old*_ group means = 1.747, 95% credible interval [1.657-1.846], Boundary_*New*_ group means = 1.642, 95% credible interval [1.558-1.730], p = 0.026) (Fig. 5b, top). Specifically, New sequences with PEs required lower threshold to arrive at a decision compared to Old sequences. In other words, a reduced threshold need to cross allows PE based memories to be recalled faster during making decisions on their temporal order.

**Fig. 5.**
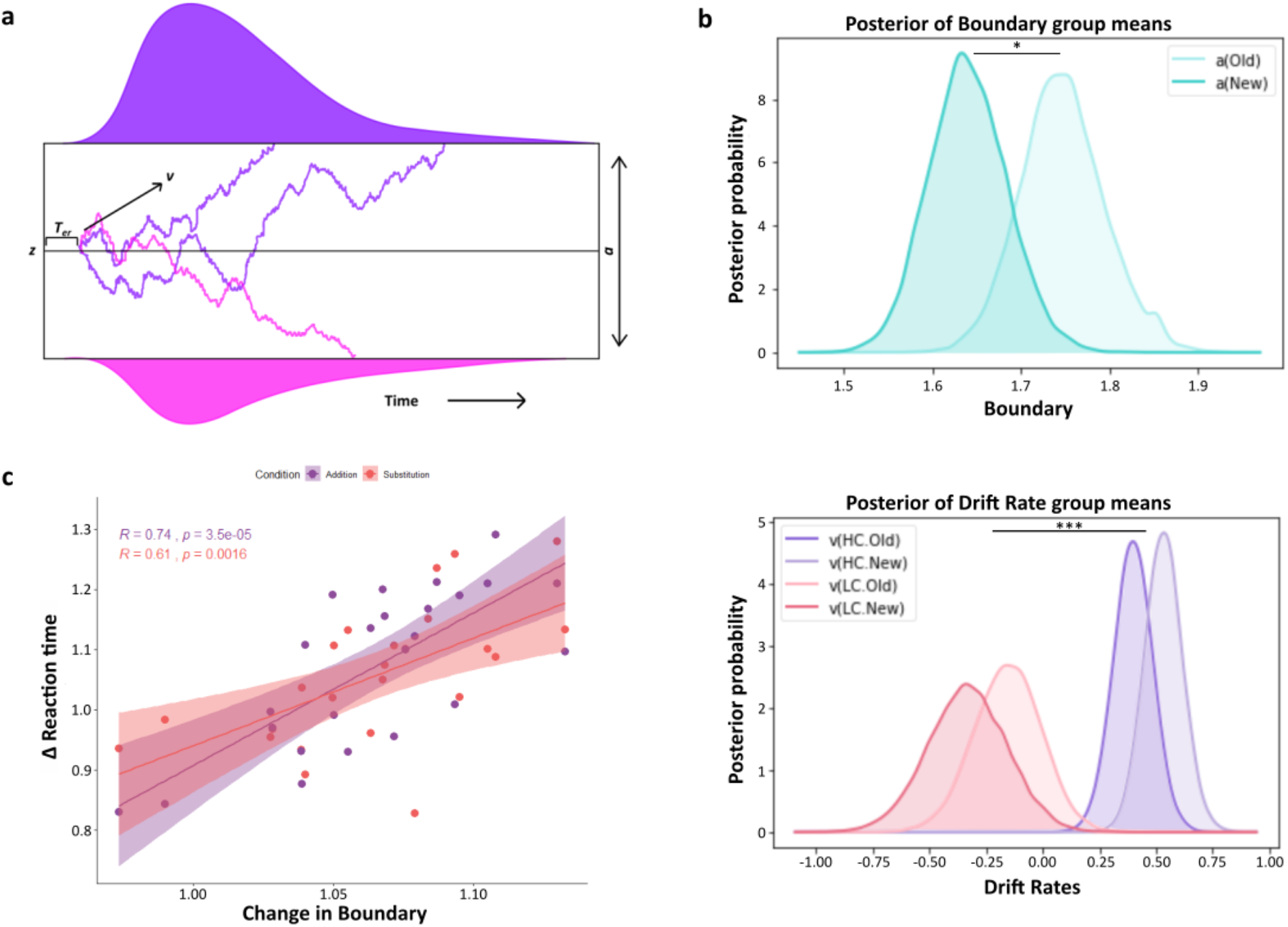
Hierarchical DDM results. **a**, An illustration of the DDM. The drift rate *v* reflects the rate of noisy accumulation of evidence till it reaches either of the two boundaries which are separated by a parameter *a*. The process starts at point *z* which may or may not have a response bias (not included in the main model) towards either boundaries, which results in the model making the response choices. The response times are a combination of diffusion process and the non-decision time *T_er_*, which includes no accumulation. **b**, Posterior density plots of the group means boundary parameter (top) showing differences (*p* = 0.026) between the Old and New sequences. New sequences were recalled faster due to the lower decision requirement while recalling compared to the Old sequences. Posterior density plots of the group means drift-rate parameter (bottom) showing no difference between the Old and New sequences but only between High and Low confidence responses (*p* <0.001). **c**, The subsequent adjustment in decision criteria made by the participants (boundary parameter) for recalling newer memories (compared to old ones) were significantly correlated with change in reaction times in Addition (Purple) *r* = 0.74, p < .001 and Substitution (Pink) *r* = 0.61, p = .0016. Dots represent individual participants.

**Fig. 6.**
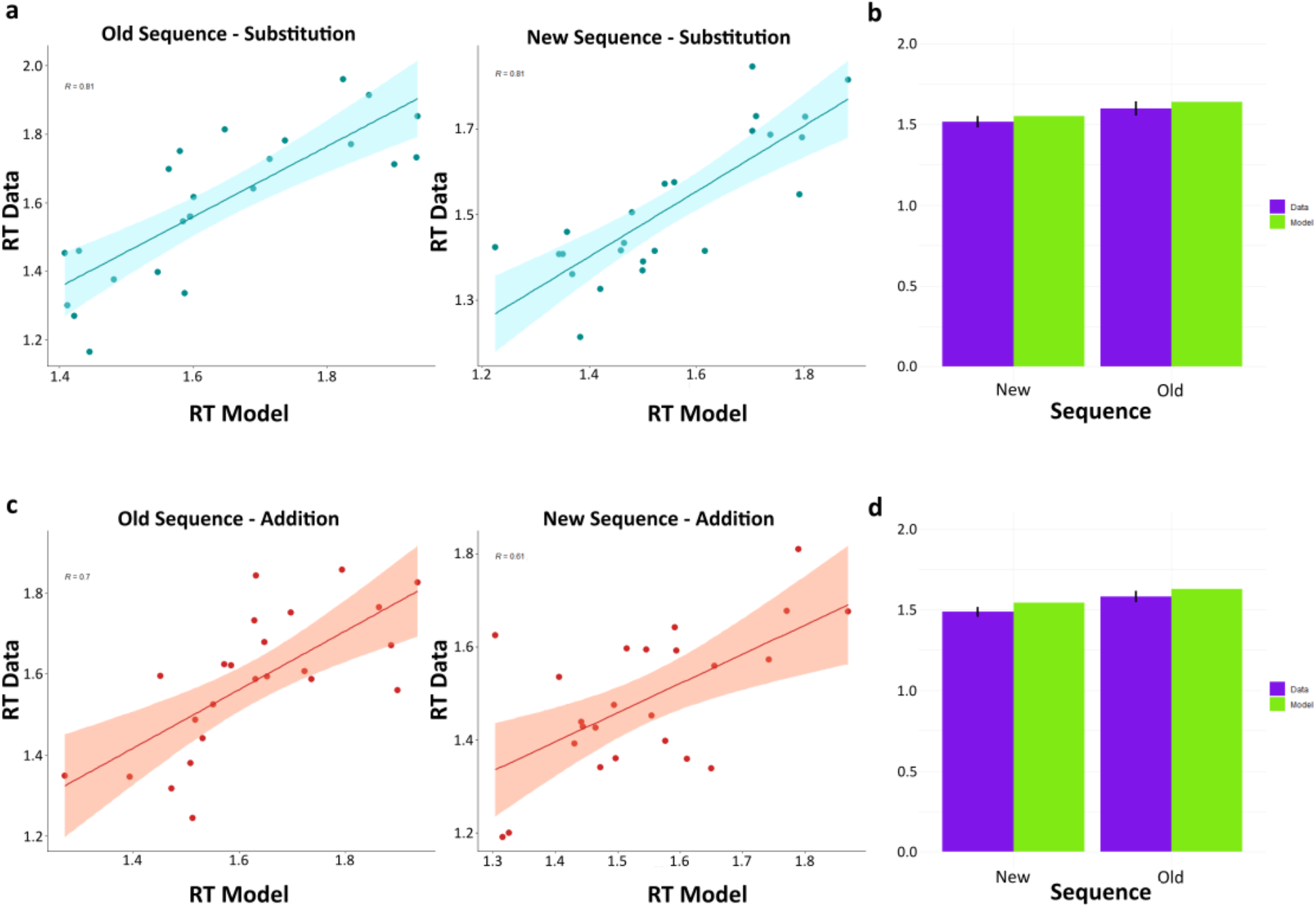
Model accurately fits the observed RT. **a**, Behavioral RT data correlates with model generated RT data in Substitution for Old sequence (left), *r* = 0.81, *p* < .00001 and New sequence (right), *r* = 0.81, *p* < .00001. **b**, Reaction time fits of model with empirical RT data in Substitution for New sequence(left) and Old sequence(right). **c**, Behavioral RT data correlates with model generated RT data in Addition for Old sequence(left), *r* = 0.61, *p* = .0014 and New sequence (right), *r* = 0.70, *p* = .00012. **d**, Reaction time fits of model with empirical RT data in Addition for New sequence(left) and Old sequence(right). Data (Purple), Model (Green). Dots represent individual participants.

We estimated the drift-rate parameter *v*, to quantify the differences between RTs in Old and New sequences. The drift-rate parameter *v,* crucially did not show any significant differences between the Old and New sequences in both Low (Drift-rate Low Confidence_*Old*_ group means = −0.153, 95% credible interval = −0.445 - 0.129, Drift-rate Low Confidence_*New*_ group means = −0.326, 95% credible interval = −0.666-0.009, p = 0.217) and High Confidence responses (Drift-rate High Confidence_*Old*_ group means = 0.389, 95% credible interval = 0.221-0.556, Drift-rate High Confidence_*New*_ group means = 0.52, 95% credible interval = 0.358-0.685, p = 0.867) (Fig. 5b, bottom). This is also in line with the behavioral result of participants’ subjective confidence judgements which did not significantly differ between Old and New sequences, in both *Substitution* and *Addition* (Supp Fig. 3). However, drift-rates did show significant differences between Low Confidence and High Confidence responses, across the Stimulus (Drift-rate High Confidence_*New*+*Old*_ group means = 0.455, Drift-rate Low Confidence_*New*+*Old*_ group means = −0.24, p < .001). This means that rate of evidence accumulation while recalling the temporal order of memory determines the subjective confidence accompanying those responses. Intuitively, a faster drift rate resulting in rapid memory activation might signal more subjective confidence in the memories.

### Change in decision boundaries correlate with change in reaction times across participants

We wanted to quantify how much of the within-subject reaction time differences between Old and New sequences were driven by the changes in decision boundaries. For this, we used the ratio of Boundary parameter *a_(Old)_*/ Boundary parameter *a*_*(New)*_ to quantify the change in evidence requirement and correlated this quantity with the ratio of RT_*(Old)*_/RT_*(New)*_ as a measure of change in reaction times for each participant. If prediction errors were affecting the decision thresholds, then there should be a correlation between how much the memories for Old sequences were weakened (as reflected by their increased reaction times compared to the New sequences) and how much the participants’ decision boundaries were increased for these sequences.

Significant correlations were found in both *Substitution* (*r* = 0.61, p = .0016) and *Addition* (*r* = 0.74, p < .001) revealing how the changes in the decision boundary drives corresponding changes in participants’ reaction times. (Fig. 5c)

## Discussion

Using episodic memory to encode events temporally helps us not only in communicating our experiences precisely to others, but also to initiate future event predictions so as to adjust behaviours accordingly. Converging evidence of the contextual and predictive functions of hippocampus, the core structure behind episodic memory, suggests a dual nature of memory recall. Here we used a movie stimulus with distinct contexts, each of which elicited a contextually specific prediction error to demonstrate this property. Our results strongly suggests updating of the temporal event structure upon encountering contextual mismatches. Specifically, the temporal order of older, inaccurately predicted sequences of memories in a given context was significantly weaker. This concomitant increase in memory strength for the newer memory sequences, was observed suggesting that PEs can selectively disrupt episodic memories in the time domain. Finally, by conceptualizing the sequence memory retrieval as an evidence accumulation process over time via a hierarchical drift-diffusion model, we offer insights into how PEs decreased evidence requirement for the new sequences during recall.

Extant literature suggests how PEs can affect declarative memory^13, 14, 15, 16^ and even destabilize it^23, 12^. But an emphasis on how temporal components of episodic memories are affected by PEs is mostly unexplored. When the predictive power of the memorized sequence of events decreased, they are weakened, leading to a reconfiguration of the temporal order such that the newer sequences are accepted as the most probable ones in that context. This was observed without an explicit reward function or multiple trial learning occurring, suggesting a mere context violation with single exposure can shape temporal memories profoundly. A very recent work^29^ which computationally modelled neuroimaging data while participants listened to temporally extended narratives demonstrated the construction and forgetting of memories were best captured by a hierarchically organized temporal context, where interaction between current and prior contexts decide memory formation. Interestingly, this was decided by a surprise or prediction error signal in the hierarchical model. Another recent study^30^ showed electrophysiological signatures of prediction error in episodic recall while participants noted a difference between an expected image and its prior encoded state. Our paradigm ensured that a prior temporal context of memories is encoded and violated at specific segments, thereby teasing out the specific behavioural effect of PEs on the sequential arrangement of memories.

Furthermore, in a second condition conceptually similar to the first, we made the participants re-experience the mispredicted segments, after inducing the PE to reactivate those sequences strongly. This enabled us to understand the interaction between reactivation, a cardinal property of memory and prediction error. Strongly reactivating memories have been shown to strengthen them^31, 32^. Strikingly, despite these Old sequences having similar accuracies compared to the New sequences, there was a stark increase in response times. In other words, even though the segments were seen again, participants were slower in recalling the temporal order signifying the fact that even re-experiencing the memories did not protect the temporal order from being affected by surprise. It could be that the memory of the rewatched segment itself wasn’t affected, rather the original temporal order in which it was encoded. In doing so, we uncovered an interaction between memory reactivation and PEs. Taken together, the main emerging finding from the above two key empirical results are a specific effect of PEs in slowing the recall of older, mispredicted memory sequences. We sought to model empirical observations to gain insight as to why this could possibly happen.

Deploying a hierarchical drift diffusion model allowed us to disentangle the mechanisms underlying these reaction time differences. That is, whether the slower reaction times were due to slower evidence accumulation rate or a higher decision threshold. Model output showed the effects of PE are due to increased decision threshold for the Old sequences, and subsequent decrease in threshold for the New sequences. Importantly, this relative change in decision threshold correlated significantly with the relative change in response times within participants. This suggests a more bottom-up, automatic decision while recalling the New sequences. We interpret this observation as participants deploying more top down control while recalling Old event sequences which reduced when they recalled the New sequences, due to the latter requiring less evidence to decide. The extent of this reduction was reflected in the speed differences observed in recalling these two sequences. In contrast, no such support was found in favour of evidence accumulation rate accounting for the effects of PEs. However, the speed of evidence build up did determine subjective confidence in the memory sequences. This confirms our behavioural finding where confidence measures did not change significantly due to the PEs, in line with similar studies^12^. Together with the behavioural results, this can explain some outstanding findings in the literature^21, 12, 33^.

The distinctive feature of reconsolidation studies is the intrusions of one set of memories when recalling another when both are learned in the same context. For example, a list of words intrudes into a second list, when both are learned separately in the same context^20, 21^. Such intrusions are by nature, asymmetric, in the sense that only the second list can intrude onto the first and never the other way around. We interpret this longstanding finding in the light of our result as follows. During remembering the temporal order of memories sharing the same context, a source confusion ensues^1, 17^ and the PE sequences owing to their lower decision threshold are recalled faster. While freely recalling memories, due to the decreased evidence required, PE memories intrudes into an older memory sharing the same context, reflecting as errors in remembering. The opposite is harder since older memories require more evidence and hence the asymmetry in temporal recall.

The discrete segmentation of continuous ongoing experiences, proposed by Event Segmentation Theory^34^, is thought to be mediated by prediction errors arising due to sudden, unpredicted contextual shifts. This effect on temporal structure can be recovered from pupillary arousal signals related marker for prediction error as shown by a recent study^35^. Such predictive principles in episodic memory are only now beginning to find common links with other prediction based learning systems in the brain^6^. Most tasks that induce prediction errors have a reward task structure learned over many trials^13, 15^. However, in our paradigm each segment created a PE only once, which was sufficient to destabilize the prior encoded sequence of events. It might be as if the accessibility of episodic memories depends on information about PEs as well while recalling, in addition to reward or value. Paradigms with both reward and non-reward, repetitive prediction errors would be required to address this. The lowered decision (criterion) requirement in the newer sequences also adds credence to predictive coding frameworks of episodic memory, where newer unexpected information is inherently prioritized to update internal models.

Alternatively, the results of our experiment can be explained with a temporal advantage effect. Particularly, since participants saw the New segments during *Day2*, they will have a recency advantage during *Day3* compared to segments encoded during *Day1*. However, we tried to minimize this recency effect by specifically instructing the participants to encode both days’ scenes with equal priority. Furthermore, our hypothesized effect of PEs on temporal order was reported in the *Addition* condition as well, which had the memory segments of *Day1* re-experienced, possibly ruling out this confound. Comparison of the New sequence memories with PEs to novel association sequences showed significant differences in memory strength and accessibility. Thus, we validated that the reported effects are due to violation of predictions only, and not due to novel associations. If the effects were purely due to association between events, then there should not have been any differences between the two sequences. But our results prove convincingly otherwise. Generally, in encoding temporal order of sequences, the association strength between two episodes depends on the informational relation on which they differ^36^. Thus, the more surprise involved, the more information gets equated with the association^37^. While we did not quantify the surprise elicited per segment mathematically, one can hypothesize the New PE sequences carry more surprise than the novel sequences following it.

One compelling question to keep in mind when understanding the effect of PEs on temporal arrangement of memories, is how it measures up with events of repeated experience. Repeated patterns help us to predict upcoming events with higher precision. In our experiment, the control sequences were repeated unchanged across both days’ viewings. Intuitively speaking, these should have better accuracies than New sequences that were seen on just once, owing to multiple exposures. Yet strikingly, the memory accuracies were similar, implying a one-shot learning for the New sequences. This is because it took these sequences only a single exposure to gain a performance similar to the repetitive sequences. Further work is needed to disentangle how the brain implements two routes of learning sequences – one-shot and repetitive learning^38^. A recent study^16^ has shown that one-shot learning is seen with PEs in paired association learning. We extend this finding to the time domain as memory sequences with PEs showed similar accuracies as control sequences.

A promising future endeavor is investigating whether such sequential reorganization of prior encoded memories can be observed in language processing as well as in navigation. Recalling, ordering and integrating memories in time is widely considered to be a function of MTL structures^10, 38, 39, 40, 41, 42^. The hippocampus, in addition to its central role in processing spatial, temporal and contextual information^43, 44^ has also been hypothesized to be a generator of sequences^45^. Future studies can uncover how contextual recall and prediction occur in this structure distinctly.

In closing, we combine the contextual and predictive properties of episodic memories, demonstrating how this mediates the temporal ordering of events. Future studies can correlate behavioural indices with neural measures in health and in disease. Since impaired temporal memory recall is one of the earliest signs of preclinical Alzheimer’s disease and mild cognitive impairment, our work can have important implications in developing aids to strengthen complex real-life memories. In addition, it also can inspire creating cognitive tools to weaken undesirable older memories in PTSD and anxiety.

## Acknowledgements

This study was supported by NBRC Core funds, Ramalingaswami Fellowships (Department of Biotechnology, Government of India) to DR (BT/RLF/Re-entry/07/2014). AB also acknowledges the support of the Centre of Excellence in Epilepsy and MEG. DR was also supported by SR/CSRI/21/2016 extramural grant from the Department of Science and Technology (DST) Ministry of Science and Technology, Government of India. AB and DR are also supported by BT/MED-III/NBRC/Flagship/Program/2019. We would also like to thank Gargi Majumdar and Dipanjan Ray for their helpful suggestions and comments on the manuscript.

## Methods

### Participants

Twenty-four healthy young right-handed adults participated in this experiment (16 males, 8 females, ages 22-35, *Mean*: 27, *SD*: 3.3). The study was approved by the Institutional Human Ethics Committee of National Brain Research Centre, India (IHEC, NBRC). All participants signed informed consent in consonance with the Declaration at Helsinki, declared normal or normal to corrected vision and no history of hearing problems, neurological and neuropsychiatric disorders.

### Materials

Participants saw 2 short films on Day 1. The selected films had engaging plots with multiple different contexts, spatially and conceptually with slightly surprising storylines. Secondly, being short films, they had no famous or otherwise identifiable actors whom the participants could easily recognize. This control on *prior* memory formation was a necessary step in our study. We wanted the natural scenes to appear as a ‘first impression’ in which aspects like characters and storyline remain unknown beforehand. Third, being an episodic sequence encoding task, it was imperative that the performance should be measured objectively. In conventional study designs with naturalistic stimuli, authors typically measure memory performance based on questionnaires, interviews etc. In order to have the same methodological rigor of controlled memory tasks we needed to edit the movie and scenes appropriately to suit our goal.

We divided the movie into different events with each event being defined by a distinct underlying context conceptually/spatially. Each event had multiple segments in them. For example, a series of segments occurring in a kitchen at night, in a bar, by the car park all constitute separate events. Each segment is defined by the actions/interactions happening between entities (people) concerning the underlying event. Thus, we divided the whole movie into different contextual events with different segments making up an event (Fig. 1). The movies were taken from YouTube, with rights for scientific purpose obtained from the creators. One was titled *The Betrayed*, about a wife finding out her husband’s affair with her best friend, with the latter trying to cover it up. Second was titled *The Man and the Thief* was about a young man helping out a girl at a railway station only to be duped by her in the end when she steals his wallet. Both movies had a combined 15 events where prediction errors occurred. Each event had 4 segments. Each segment was 7s long, with a 1s interstimulus interval, where a black blank screen is presented between every segment. This also allowed us to reorient participants’ attention between the scenes.

#### Experimental procedure

##### Day 1: Formation of Priors

All participants saw both movies during *Day1* prior formation. The order of the movies seen was counterbalanced across participants. Each event had 4 segments on *Day1,* for both movies. They were named (in order of their appearance in the event) *Start*, *PrePE*, *Old* and *PostPE*. Each event began with the *Start* segment.

##### Day 2: Inducing Prediction Errors

To induce PEs, the original versions of the movie were edited into 3 versions. First version being the *Prior* was shown to all the participants on *Day1*. The second version was the *Substitution* version with a segment substituting another within an event and the third version was the *Addition* version with an additional segment inserted in an event. These scenes termed *New* segments were contextually fitting so as to avoid making abrupt transitions while watching. Importantly, because of this, we could reliably introduce contextual PEs in both movies. Furthermore, on *Day2*, participants saw the two movies in either of these two versions.

First segment of every event was termed as *Start* since it was always the first event to be shown in a segment and was unchanged across both days’ viewings (for this reason, it acted as Control). This also enabled the subjects to predict the upcoming segments in that specific event on *Day2*. Second segment was termed the Pre-Prediction Error segment (*PrePE* henceforth) since the prediction violation segment always happened after this. This segment was also similar to a Control in that it did not deviate from prior viewing as well. The segment that followed *PrePE* was termed as *Old* because this segment was shown on *Day1* but was replaced on *Day2* in the *Substitution* condition. The segment replacing it on *Day2*, the prediction error segment, was labelled as *New*. The segment that occurred in the event right after the *New* segment was termed Post-Prediction Error (*PostPE* henceforth) because it always followed the Prediction error segment.

In the *Addition* condition, *Start* and *PrePE* remained the same, similar to *Substitution*. The Prediction error segment, *New*, occurs after *PrePE*. The *Old* segment as seen on *Day1* which was expected to happen after *PrePE*, was seen after the *New* event hence termed aptly as *PostPE* in this condition.

Thus, in the *Substitution* condition, on *Day2* each event had a *Start* segment, followed by the *PrePE* segment, then the *New* segment inducing prediction error (since subjects were predicting *Old*), and afterwards *PostPE* segment. In the *Addition* condition, the *Old* (which was seen after the *New*), functioned as the *PostPE* segment. We purposely put the PE segments in the middle of events, rather than beginning or end to control for primacy and recency effects, respectively.

*Betrayed* was 5mins 42s in both *Prior* formation and *Substitution* condition, and 6mins 55s in *Addition* condition. It had 11 different events of which 9 were altered during second days viewing. *The Man and the Thief* had 6 contexts instead with a duration of 3mins 14s in Prior and *Substitution* condition, and 4mins 2s in *Addition*. All the 9 events had 4 segments (termed from start to end: Start, *PrePE*, *Old*, *PostPE*) during Prior viewing. *Old* segment gets replaced by *New* in *Substitution* while in *Addition, New* is inserted in between *PrePE* and *Old* incurring a total 5 episodes in one contextual event in this condition. Hence in *Addition*, we deliberately took out a segment from each event in the Prior version only to be added back in *Day2* so as to induce the contextual prediction errors.

Participants were instructed to pay full attention to both Days’ movies and not to use any specific strategies of encoding. They were asked on *Day2* if they noticed changes in scenes compared to the previous day to confirm for attention for both days’ viewings, but were not asked to recall them in any way.

##### Day 3: Sequence Memory Test

On *Day3* participants underwent a sequence memory test. The Sequence Memory test was a two-alternative forced task choice (2AFC) task. A screenshot from each segment was shown besides a screenshot of an adjacent segment within an event and subjects were asked to choose the first one that came in the movie. The stimulus presentation on *Day3* was using PsychoPy software (46).

It was designed in such a way that segments exclusive to one day (such as *New* and *Old* in *Substitution* condition) were not seen together to prevent a conflict of decision. Since participants were instructed to encode both Days’ movies equally without giving preference to one, there were no discrepancies between choices.

Screenshots of adjacent segments within an event were shown side by side (in normal and reversed order randomly) and participants were asked to indicate which one came first in the sequence. This enabled us to probe into the sequential association strength between those two segments, within an event. We had categorized the association strength between the *Start* segment and *Pre-PE* segment within an event as Control sequence. *PrePE* segment and *Old* segment (*PrePE*-*Old*) which reflected the strength of the older (*Prior)* sequence was termed Old sequence. *PrePE* segment to *New* segment (*PrePE*-*New*) was called New sequence and finally the sequence of *New* segment to *PostPE* (*New*-*PostPE*) segment as Novel association sequence. Trials were binned across participants into these categories by movies and conditions for analysis.

Trials were shown for 2500ms with 500ms ITI between them. Subjects were asked to answer as accurately and fast as possible. After the response subjects had to choose their confidence on a 3-point scale - High Confidence, Low Confidence and Guess. There were 180 trials in total per participant. We calculated the accuracy by including only the High Confidence and Low Confidence hits; Guesses were completely excluded from analysis. Confidence performance of High Confidence responses were derived by counting the percentage total of High Confidence hits per total hits (High Confidence + Low Confidence).

### Behavioral Data Analysis of Sequence memory

In order to determine the sequence memory performance, we categorized the trials into *New* sequences (temporal order judgements between *PrePE* and *New* segments), and *Old* sequences (temporal order judgements between *PrePE* and *Old* segments). All trials were binned into either *Addition* or *Substitution* by participants depending on which version of the movie they watched on *Day2*. Furthermore, we also categorized trials into Novel sequences (temporal order judgements between *New* and *PostPE* segments) in *Substitution* and *Addition* as well.

Subjective confidence in memories is usually teased out via either numerical scales (1-10) or nominal scales (sure, not sure). Here we deployed a version of nominal scale having High confidence, Low confidence and Guess as labels in a 3-point scale to test the strength of subjective opinion. We calculated and compared the percentage number of High Confidence responses given by participants out of the total (High Confidence + Low Confidence) responses in Old and New sequences.

### Statistical Analysis

In addition to the normal statistical tests, we also did a Bayes factor (BF) analysis on the paired *t-tests*^47^ using a default Jeffreys Zellner Siow (JZS) prior using the *BayesFactor* package in R.

### Hierarchical Drift-Diffusion Model

We used a Hierarchical version of the DDM to fit the behavioral data^48^. The DDM^49^ is one of a general class of sequential sampling models aptly suited for reaction time data from two-alternative forced choice paradigms^50^. It operates under the assumption that decisions are made by accumulated evidence from a noisy sensory signal. The decision is made when the evidence crosses a threshold. The main parameters in DDM is a drift rate, a rate of accumulation of evidence and the thresholds of the boundaries for these to cross or evidence required for the decision to be made. Additional components include a response bias, which determines whether the responses towards either boundaries are biased or not depending on the starting position. The total reaction time is assumed to be a combination of processing time required to make that decision, as well as encoding time of the stimulus and the time required for the motor execution response. The latter two are assumed not to vary, and are combined as another component called non-decision time. Thus, the DDM gives the response choice depending on which boundary threshold (upper or lower) is crossed and gives the response time as a combination of the total time required to cross these boundaries and the non-decision time involved^49, 50^.

The HDDM toolbox^48^ in python was used to model the data. Bayesian inference which naturally fits a hierarchical DDM, can be used to not only recover the parameters but more importantly to estimate the uncertainty involved in the model parameters. They also provide solutions for parameter estimations of individual participants which are assumed to be drawn from a group-level prior distribution and are constrained by it. Furthermore, Bayesian methods are more powerful when the trial numbers are low which is desirable since the usual DDM requires larger datasets. The joint posteriors of all the model parameters are estimated by standard Markov Chain Monte Carlo (MCMC) methods^51^. A direct Bayesian inference was performed on the posteriors of different conditions by computing the overlap between their distributions.

In our main model, we allowed both drift rate and boundary parameters to vary with Stimulus, that is *Old* and *New* memory sequences. This was to compare which of the two parameters demonstrated the empirical effect of PEs, i.e., reaction time. Drift rate also varied across Confidence. Confidence here being High Confidence and Low Confidence as Guesses were discarded from main analysis. Non-decision time parameter was set to vary across *Substitution* and *Addition*. We also generated other models with different combinations of drift rate and boundary varying with Stimulus and Confidence. Models were adjudicated using Deviance Information Criterion (DIC) which penalizes *Addition*al model complexity^52^. DIC values above 10 are generally considered significant, with the lowest having best goodness-of-fit. (Supplementary Fig. 4, Supplementary Table 1)

For each model, we used MCMC methods to generate 100,000 samples from the posterior distribution and discarded the first 20,000 samples as burn-in. After visually inspecting the chains and autocorrelation for proper convergence, the Gelman Rubin R-hat statistic was confirmed to be between 98-1.02^53^. The latter was computed by running the (main) model 3 times, and checking within-chain and between-chain variance. All procedures were done as described in the HDDM toolbox in python^48^.

